# Detection of insect health with deep learning on near-infrared sensor data

**DOI:** 10.1101/2021.11.15.468635

**Authors:** Emily Bick, Sam Edwards, Henrik H. De Fine Licht

**Author notes:** These authors contributed equally to this study (Emily Bick, Sam Edwards).

## Abstract

Conventional monitoring methods for disease vectors, pollinators or agricultural pests require time-consuming trapping and identification of individual insects. Automated optical sensors that detect backscattered near-infrared modulations created by flying insects are increasingly used to identify and count live insects, but do not inform about the health status of individual insects. Here we show that deep learning in trained convolutional neural networks in conjunction with sensors is a promising emerging method to detect infected insects. Health status was correctly determined in 85.6% of cases as early as two days post infection with a fungal pathogen. The ability to monitor insect health in real-time potentially has wide-reaching implications for preserving pollinator biodiversity and the rapid assessment of disease carrying individuals in vector populations.

**One sentence summary:** Automated optical sensors distinguish between fungus-infected and healthy insects.

## Main text

Ecosystem services provided by insects such as pollination or predatory insect pest control are contingent on having healthy insects. The health of insects is threatened by chemicals (*1, 2*), and microbial pathogens (*3, 4*), especially insect-pathogenic fungi, which are important regulators of natural insect populations (*5, 6*). In addition, warming temperatures and human mediated radiation and chemical exposure may further exaggerate the negative influence on insect health (*7, 8*), and there is thus a dire need for monitoring and studying insect health to preserve insect biodiversity.

Insect disease is typically identified by visual diagnosis of diseased insects post-mortem, or via molecular screening with probes or primers (*9*). These methods are time consuming and usually only cover a small fraction of entire insect populations. Optical insect sensors have recently become more widely accessible by significantly reducing the physical size and exchanging lasers with eye-safe LED lights (*10*). Automated insect monitoring sensors are currently used to remotely count insect flights (*11*), record wing beats (*12*), and classify insects to species (*13, 14*). The incorporation of optical signals with machine learning algorithms enables increased accuracy in remote insect species classification (*13*). Here we used autonomous near-infrared sensors to study if backscattered light and wingbeat frequencies can differentiate between infected and healthy insects.

To establish age-controlled cohorts of healthy and infected insects, we used the common house fly (*Musca domestica*), which is a globally distributed insect that can act as a vector of >100 human disease-causing microbes (*15*). Virgin house fly males were infected with the obligate fungal pathogen *Entomophthora muscae* (*16, 17*). This fungus has an incubation period of 6-7 days during which it initially grows exponentially inside infected flies (*18*), before it takes over the behavior and kills its infected host. Fifty healthy or infected five day old male flies were released into a flight cage equipped with an autonomous near-infrared sensor (*10*) with a measurement volume of 17.5 L and the angle between the emitter and receiver (*θ* s) set at 18 degrees (fig. S1). Fly cohorts were continuously fed inside cages and flight events monitored for >7 days, which resulted in >10,000 recorded flight-events per day.

We used a Convolutional Neural Network (CNN) approach (*19*) to train the network on 8,000 events per day of light back-scattered by the flying insects with a resulting classification as either ‘infected’ or ‘control’ (fig. S2). For each day, 1,000 flight-events were used as validation set and the process was refined over 20 epochs before the final model was tested on 1,000 flight-events (fig. 1). Two days post exposure to infectious fungal spores, the day-2 model was able to correctly predict infected insects in 61% of cases (fig. 1). As the disease progresses until host death at 6-7 days post infection, the applicability of discerning infected from healthy flies increases from 61% to 99% (fig. 1). This is in concordance with the gradual colonization of the hemolymph and insect body by fungal cells from the initial exposure (*18*), and thus correlates with disease progression.

**Fig. 1.**
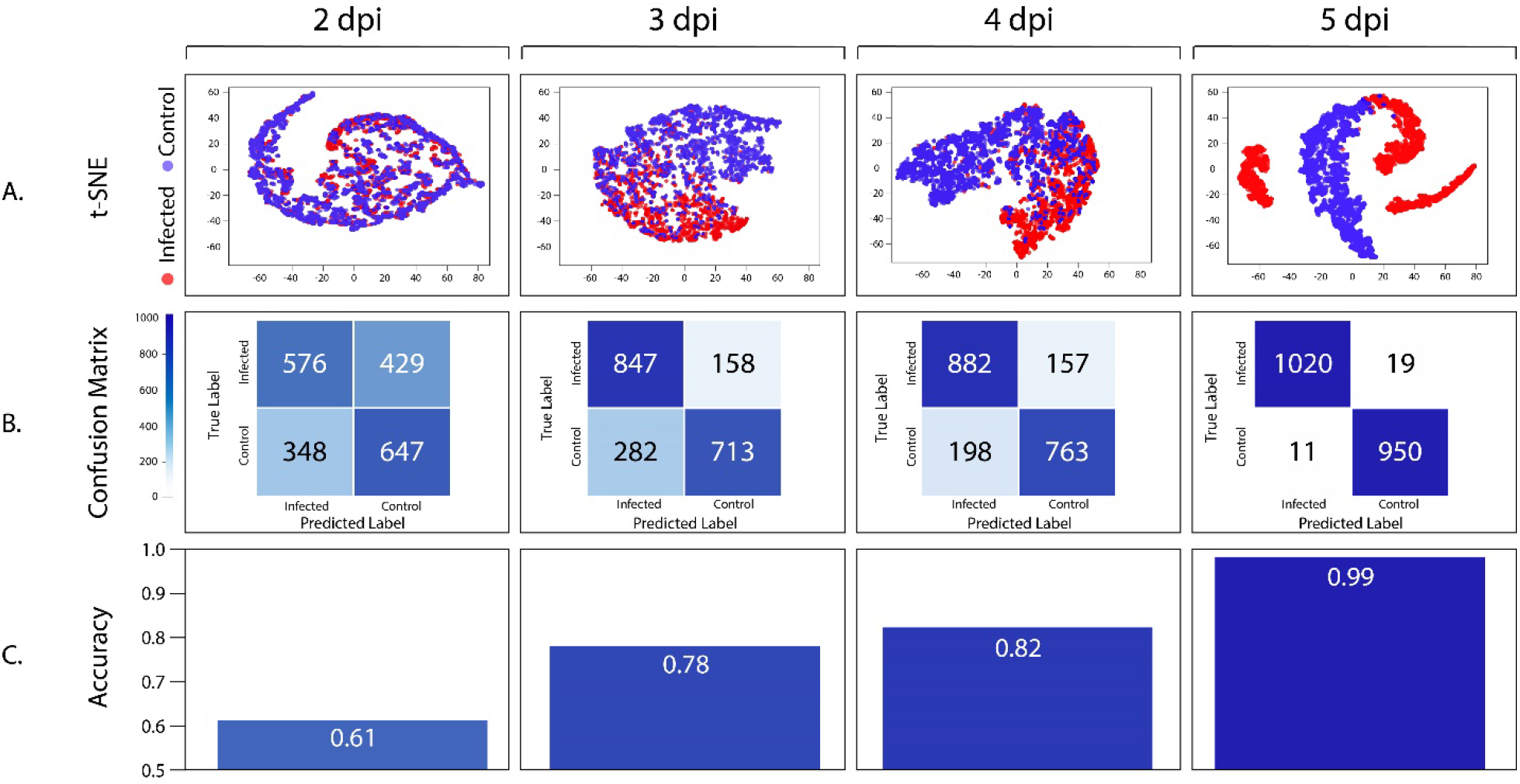
Evaluation of Convolutional Neural Network models trained on back-scattered light from healthy and infected house flies. Evaluation of models trained on data from 2-, 3-, 4-, and 5-days post infection (dpi) with **A. A** t-distributed Stochastic Neighbor Embedding (t-SNE) plot, **B**. A Confusion Matrix of true label versus predicted, and **C**. Accuracy of model predictions.

Next, we sought to use the CNN model with the highest accuracy on a new dataset with flies at a different age and in various stages of infection. Data from 50 flies infected at nine days old, released into the flight cage at ten days old, and recorded until age 16 days (note the average lifespan of house flies are three-four weeks (*20*)) were obtained as described before and analyzed with our model trained with the five days post infection data (fig. 1). This showed that infrared light back-scattered by flying insects in combination with an unsupervised deep learning algorithm are able to correctly predict infected insects in ca. 79.0% of cases as early as two days post infection (fig. 2). Infection with *E. muscae* in house flies can normally only be determined two days post infection using molecular or microscopic diagnostic procedures (*18*), and behavioral effects on infected hosts are normally not observed.

**Fig. 2.**
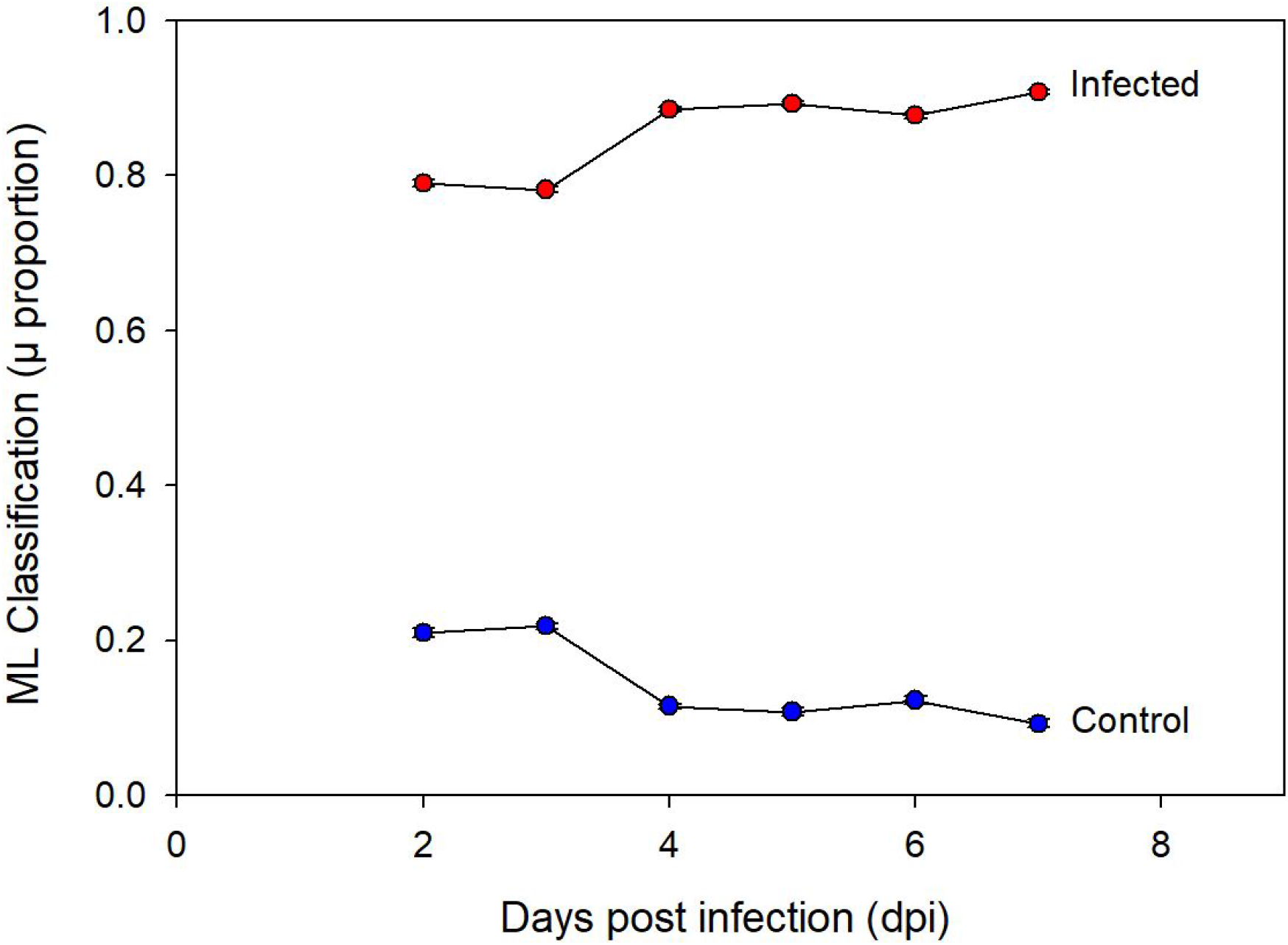
Application of the most accurate model trained on 5 days post infection (dpi) events was applied to a new dataset of flies infected at age 9. The model predicts a likelihood (proportion) that an event falls in the ‘infected’ category and a separate likelihood (proportion) that it falls in the ‘control’ category. The mean (μ) ± SE of likelihoods predicted by the model are reported for infected flies (red) and ‘control’ flies (blue).

We also investigated average wing beat frequency, which differed significantly between control and infected treatments (ANOVA: P < 0.001) (table S1), but showed overlapping variation between healthy (170.25 Hz, SD = ± 20.67) and infected (day 5: 183.03, SD = ± 20.81; day 10: 171.47, SD = ± 20.11) flies (fig. 3, table S2). Same-sex sexual behavior and aggression (*21*) resulting in damaged wings may partly explain some of the variation in recorded wing beat frequencies in both infected and healthy flies. While average wing beat frequency did not discern between healthy and infected flies across all ages, the total number of recorded flights were lower in infected flies compared to controls (ANOVA: P = 0.0177) (fig. S3, table S1), and the maximum recorded wing beat frequencies in flies infected at age 5 and 10 were higher (273.76 Hz and 274.52 Hz, respectively) compared to healthy control flies (247.07 Hz).

**Fig. 3.**
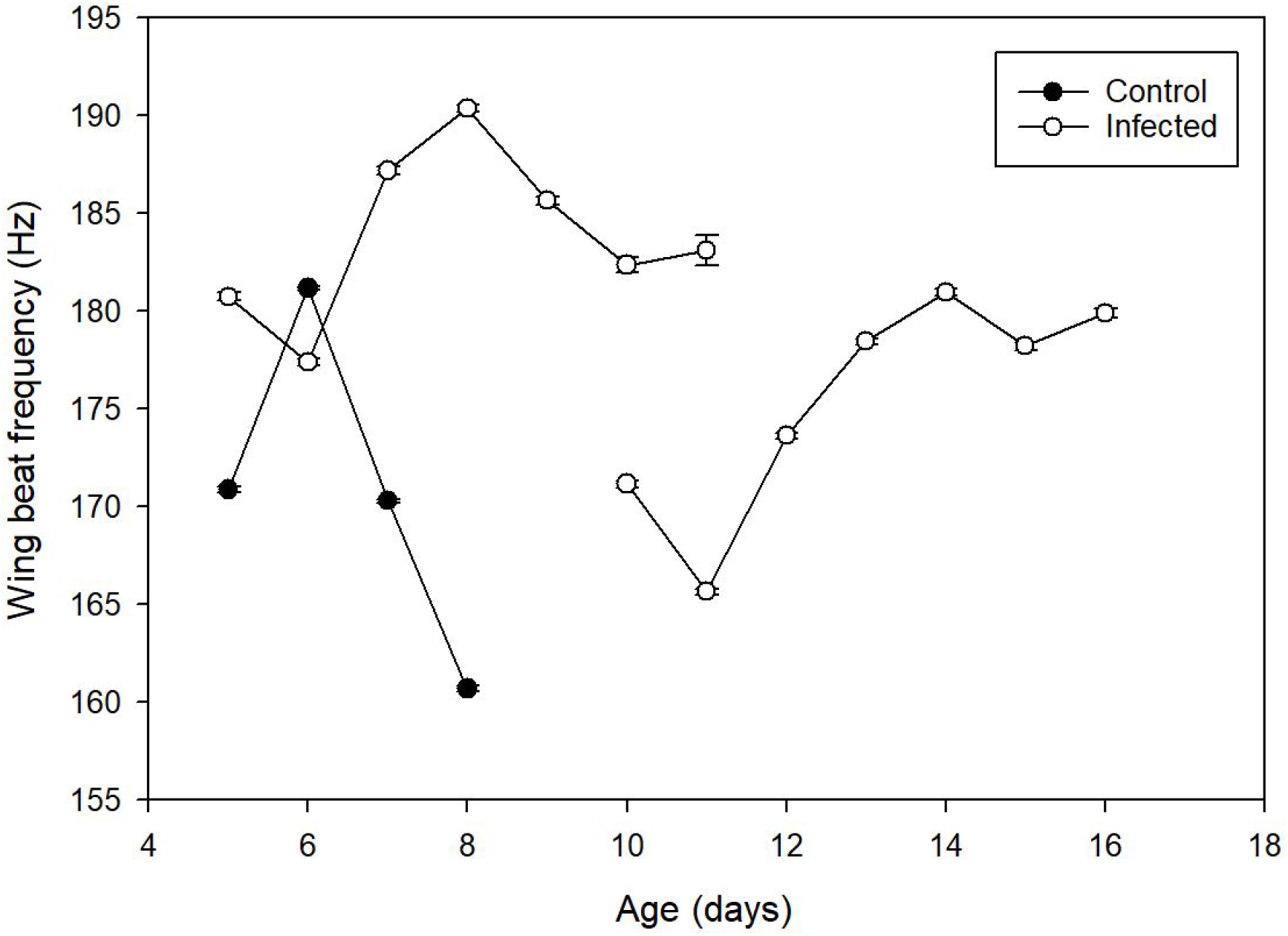
Variation in wing beat frequency with age in infected and control flies. Infected individuals were observed from the second day post infection through the eighth day post infection.

Our results provide the first evidence that near-infrared optical LED sensor technology in combination with deep learning algorithms can be used to non-destructively differentiate healthy and infected flying insects from less than 0.36 seconds of flight time. The ability of the trained CNN model to detect infected individuals with 85.6% mean accuracy as early as two days post infection where less than 100 fungal cells may be found inside infected flies (*18*), suggests that other types of viral, bacterial or parasitic insect diseases can potentially be detected. While the differences in backscattered light used by the CNN model to differentiate between healthy and infected are unknown, the model certainly responds to changes in flying behavior. For example, vertically inherited *Wolbachia* bacterial endosymbionts are known to change searching behavior of infected mosquito hosts (*22*), which opens the possibility for using backscattered light to determine *Wolbachia* infection status. While work is needed to test the classifier under field conditions in combination with conventional validation methods, the ability to rapidly detect the health status of insects solely based on back-scattered light from flying insects is a promising technology with potentially wide-reaching implications in several areas. First, the ability to non-destructively monitor insect health could help rapidly inform efforts to preserve pollinator biodiversity. Second, when chemicals or microbial pathogens are used against insect pests, LED or laser assisted monitoring could rapidly inform on the efficacy of the treatment. Third, for mosquito-borne human diseases, such as Malaria (*23*) and Dengue fever (*24*), the proportion of disease-transmitting insects could potentially be directly monitored and identified using the methodology described here.

## Acknowledgements

Thanks to Dr. Freja Thoresen, Dr. Marta Montoro, Rami El Rashid, and Klas Rydhmer from FaunaPhotonics ApS. for assistance with the data acquisition and analysis. Thanks to Alfred Strand for assistance with estimating measurement volume and to Laurence Still for editing and interpretation. Thanks to Dr. Jesper Lemmich and FaunaPhotonics ApS. for coordinating resource support for this work.

## Author contributions

Conceptualization: EB, SE, and HHDFL. Methodology: EB, SE, and HHDFL. Investigation: EB and SE. Data analysis: EB and SE. Funding acquisition: EB and HHDFL. Writing – original draft, review and editing: EB, SE, and HHDFL.

## Data materials and availability

Datasets referred to in the manuscript are available upon reasonable request.

## Supplementary Material

### Materials and methods

#### Insect model

We used virgin male house flies, *Musca domestica* (wt strain 772a), which were provided by the University of Aarhus, Denmark, as pupae. Flies were sexed and sorted within 24 hours of emergence as adults from pupae and kept in groups of ten to minimize damage to the wings caused by aggression and same-sex sexual behaviors. The flies were fed with ad libitum food (1:1 ratio caster sugar:semi skimmed milk powder) and demineralized water using a 10ml vial plugged with cotton.

For infected treatments, three house fly cadavers newly killed (within six hours of death) by *Entomophthora muscae* (isolate no.: KVL21-01, University of Copenhagen, Section for Organismal Biology Entomopathogenic fungus culture collection) were used to infect ten healthy male house flies. The three cadavers used, with a minimum of one male and one female, were fixed head first in 5% water agar in medicine cups. Ten adult male house flies were anaesthetized at 5°C and placed with the cadavers for 24 hours in a saturated humid chamber. The cups were perforated for aeration and placed upside down to allow for spore showers by the infected cadavers. For the healthy treatment, mock-infections were performed in the same manner as for infected treatment but using uninfected dead house flies, which were euthanized by five-minute exposure to −18°C. All flies used in the experiment showed no developmental deformities to the body or wings and appeared healthy prior to being used for experimentation.

#### Cohorts

Three cohorts, each composed of 50 flies, were experimentally released into a label cage (see Data collection section below for label cage specifications). The first cohort, termed ‘control’, was mock-infected four days post-pupation and released in a label cage aged five days. These 50 healthy flies were kept in the label cage for 96 hours (four days – from dpi 2 to 5) before being removed. The second cohort was made up of 50 flies the same age as cohort one but were infected with *Entomophthora muscae*. The second cohort remained in the label cage for nine days. The third cohort were infected nine days after emergence as adults from pupae and released aged ten into the label cage and again remained there for nine days.

The labelling of days-post infection (dpi) is defined as follows: Day 1 (0-24 hours) refers to the time of infection with cadavers; Day 2 (24 to 48 hours) refers to the day of release into the label cage and the beginning of sensor monitoring; Day 3 (48 to 72 hours), Day 4 (72 to 96 hours) and Day 5 (96 to 120 hours) are days of monitoring of flight.

#### Data collection

Data collection occurred at the University of Copenhagen, in Taastrup, Denmark from 05.19.2021 to 06.08.2021. An autonomous near-infrared sensor (detailed description of the sensor is provided in (*1*), produced by FaunaPhotonics ApS., Copenhagen SV, Denmark) monitored each cohort of male *Musca domestica* flies. Specifically, we used a 17.5 L measurement volume of the sensor, as the angle between the emitter and receiver (*θ*s) was set at 18 degrees. The flight recordings occurred in a neoprene and mesh cage (termed ‘label cage’) that was >20 times the sensor’s measurement volume, to allow for sensing of typical flights.

#### Machine learning algorithm

For each insect flight, backscattered light is recorded as a 1-dimensional time series of backscattered light intensity, hereafter called a flight-event. A single flight may result in up to eight flight-events, as that flight may be recorded on one to eight channels (2 wavelengths x 4 photodiodes). Each event used is between 240 to 1500 samples in length. If a selected event is longer than 1500 samples, the middle of the event was used. So, each individual event ranges from 0.0576 to 0.3600 seconds long.

Four machine learning algorithms were trained, validated, and tested against the first two experiments (control and young infected). Each model was trained on a single day’s flight data, from two through five days post infection. For each algorithm, 10,000 random events per group were fed into an 8-layer Convolutional Neural Network (CNN) implemented in Python using Keras (*2*) (fig. S2). Events were separated into 80% training, 10% validation, and 10% test sets. The CNN uses four convolutional layers which use the ReLU activation function, two pooling layers which use the MaxPooling1D and GlobalPooling1D functions, and a dropout rate of 0.5 on the penultimate layer. The output layer used a softmax activation function to categorize events as ‘infected’ or ‘control’. The model’s application to the test set was visualized with a t-SNE plot, evaluated with a confusion matrix, and the proportion accuracy reported (fig. 1).

The machine learning model with the highest accuracy was applied to the entire dataset of events in the second and third cohort (5 days post infection and 10 days post infection), excluding the day used to train and evaluate the model to avoid data leakage (fig. 2). The model predicts a likelihood (proportion) that an event is from an infected fly or a control fly. We report the output of the model as a mean and SE of these two likelihoods.

#### Wing beat frequency and flights

Wing beat frequency (WBF) is calculated from each event (*1*). A random subsample of 1000 WBF events per treatment was used for analysis to standardize sample size across experiments. The subsample covered 250 events per day of interest, *viz*. 2-, 3-, 4- and 5-days post infection, for each of the three experiments. This sample size was selected for consistency with the test set size. Mean flight number and WBF comparisons were performed using one-way analyses of variance (ANOVAs) to determine statistical differences between cohorts (table S1). The one-way ANOVA was selected for comparison of means between a quantitative dependent variable, such as WBF or flight number, and a independent variable, such as experiment. Tukey’s honestly significant difference (HSD) post hoc analyses were used with the above ANOVAs to determine statistical differences with multiple comparisons between cohorts (table S2). All flight numbers and wing beat frequency analyses were performed using R (version 4.0.2) (*3*).

## Supplementary tables and figures

**Table S1.** Results of one-way analysis of variance (ANOVA) on wing beat frequencies between infected and control flies from the three cohorts of 50 flies.

| Cohort comparison | Data compared | Sum Sq | Mean Sq | F value | P value |
| --- | --- | --- | --- | --- | --- |
| All cohorts | Flight numbers | 7.592e+08 | 379613643 | 5.162 | 0.0177 ** |
| All cohorts | Subsamples<br>WBF | 99508 | 49754 | 118 | < 0.001 *** |

**Table S2.** Results of one-way analysis of variance (ANOVA) with Tukey’s honestly significant difference (HSD) post hoc analyses comparing means between cohorts.

| Cohort comparison | Flight number<br>P value | Subsample WBF<br>P value |
| --- | --- | --- |
| Infected_age4 vs control | 0.0153136 * | < 0.001 *** |
| Infected_age9 vs control | 0.0692365 . | 0.3769392 |
| Infected_age9 vs infected_age4 | 0.6334101 | < 0.001 *** |

**Fig. S1.**
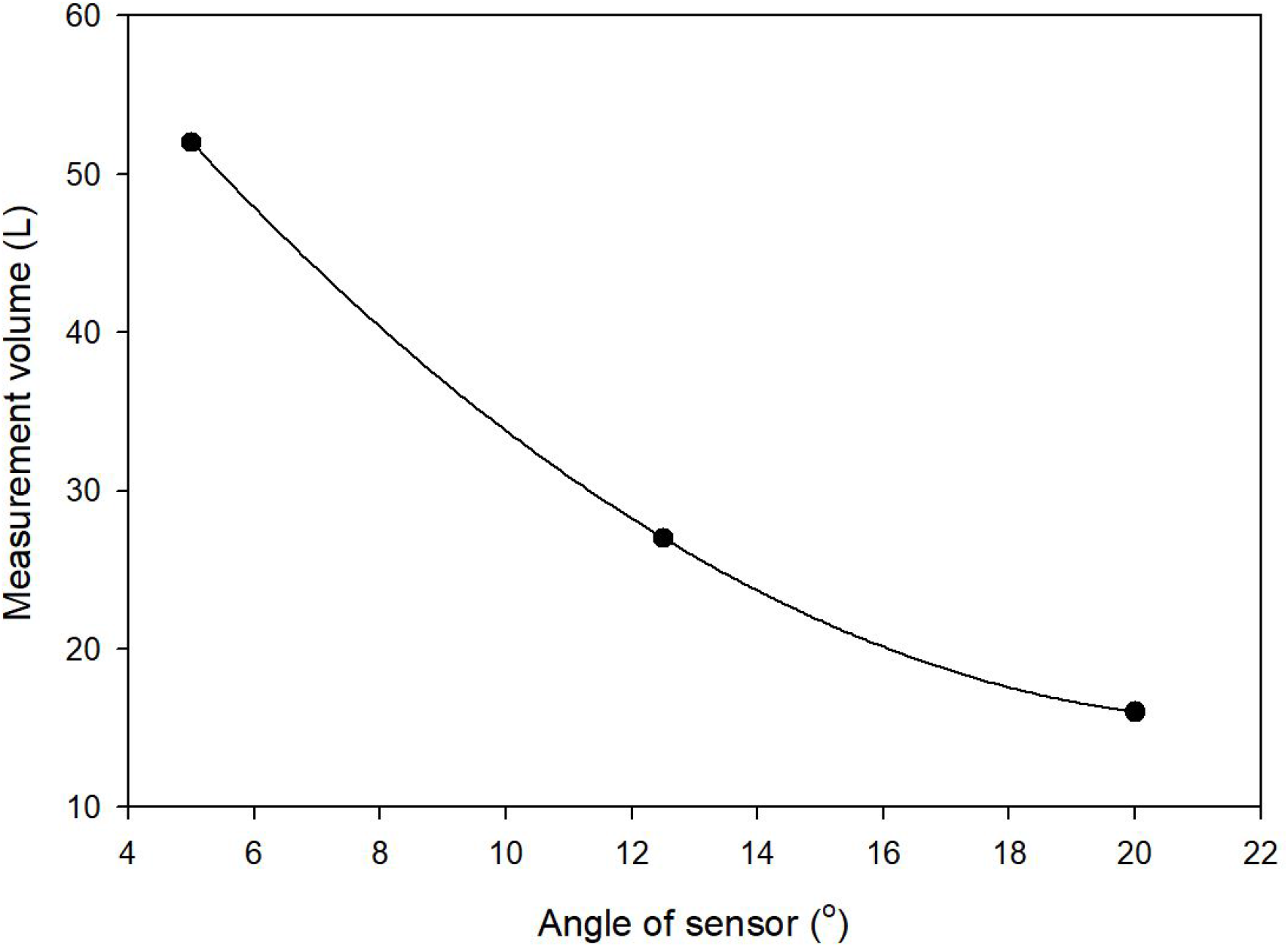
To determine the measurement volume of the sensor based on an untested angle between sensor emitter and receiver, we fit a quadratic equation to data relating angle of sensor to measurement volume. We then solved the equation for the known sensor angle.

**Fig. S2.**
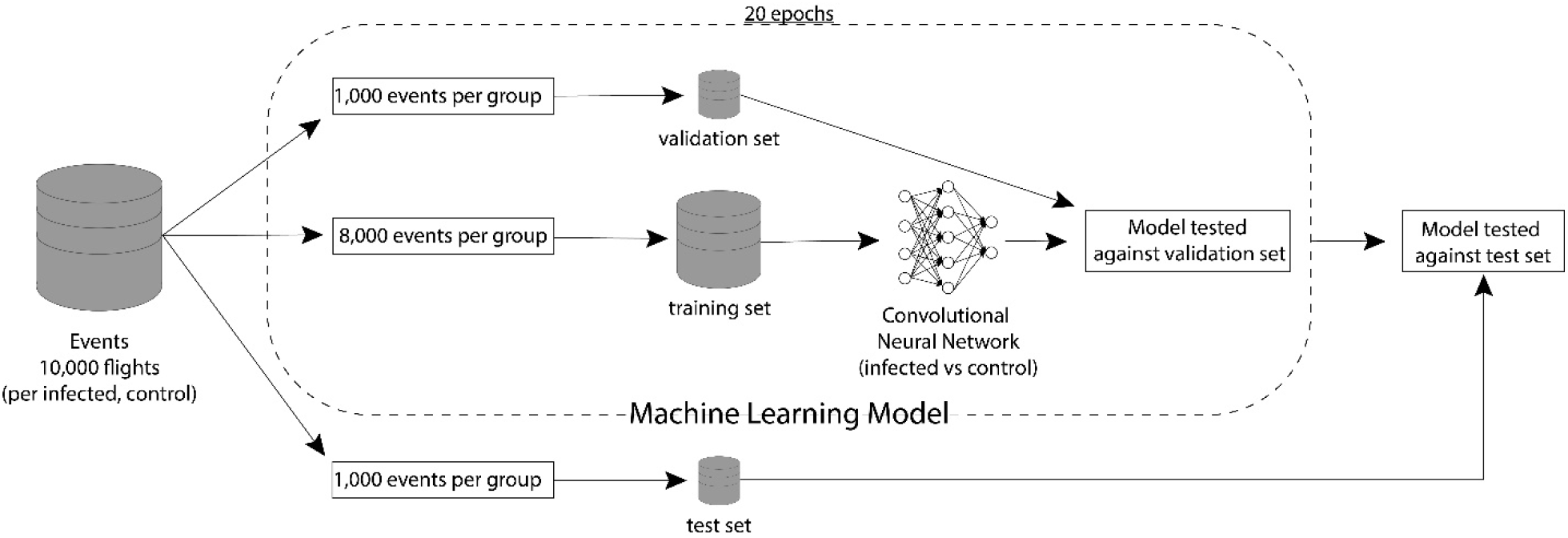
Diagram of construction and testing of the machine learning model. 10,000 events per group (infected, control) were randomly selected. Events were split into the validation set (1,000 per group), training set (8,000 per group) and test set (1,000 per group). A Convolutional Neural Network was used to classify events as infected and control, and was refined over 20 epochs. Validation set was used to monitor the CNN’s progress. We report accuracy of classifier on the test data.

**Fig. S3.**
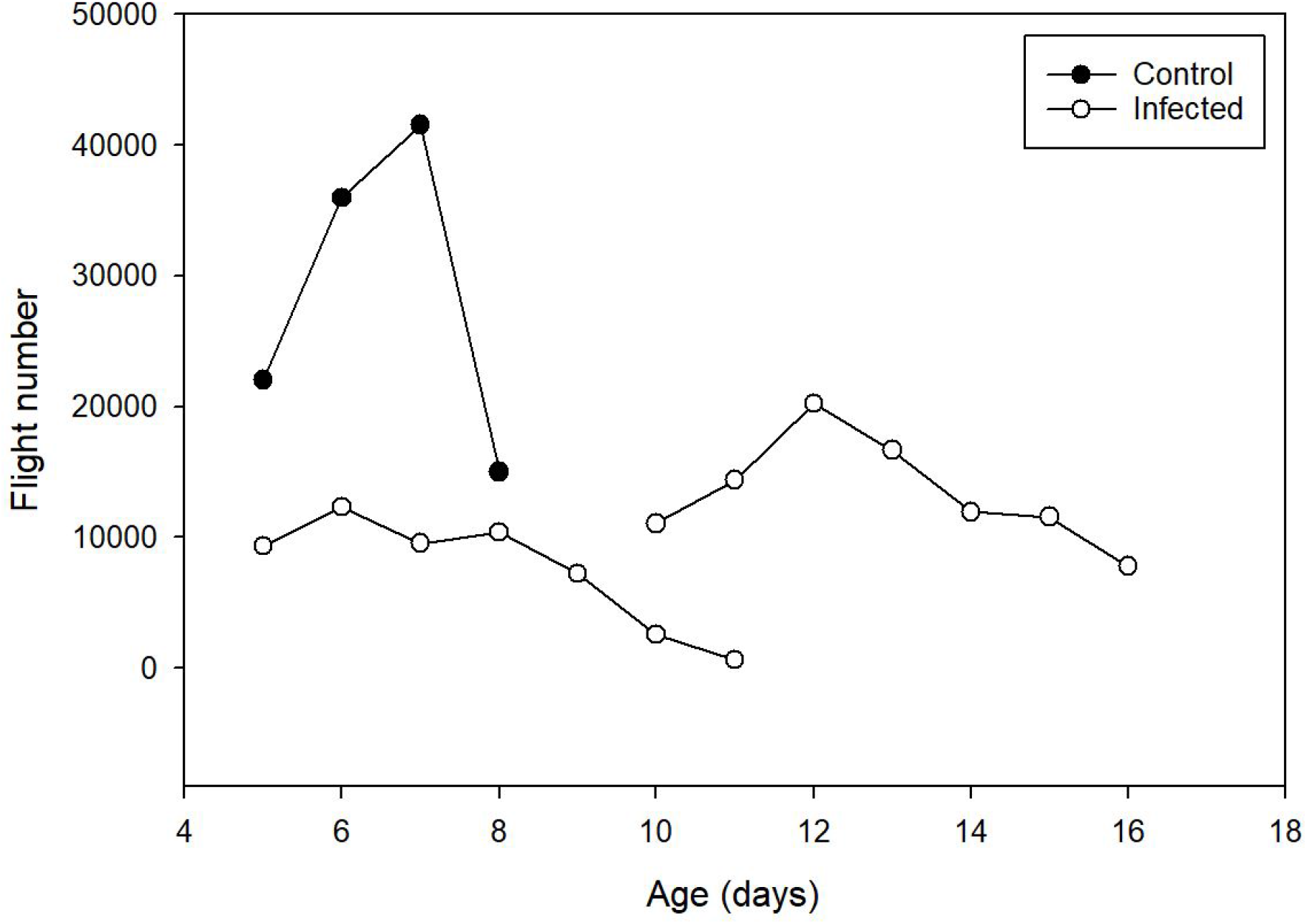
Insect activity was tracked for each experiment by counting flights per day for each experiment.

## References

1. R. J. Gill, O. Ramos-Rodriguez, N. E. Raine, Combined pesticide exposure severely affects individual-and colony-level traits in bees. Nature. 491, 105–108 (2012).

2. N. Tsvetkov, O. Samson-Robert, K. Sood, H. S. Patel, D. A. Malena, P. H. Gajiwala, P. Maciukiewicz, V. Fournier, A. Zayed, Chronic exposure to neonicotinoids reduces honey bee health near corn crops. Science. 356, 1395–1397 (2017).

3. A. Vilcinskas, Pathogens associated with invasive or introduced insects threaten the health and diversity of native species. Current Opinion in Insect Science. 33, 43–48 (2019).

4. B. Lovett, R. J. S. T. Leger, in The Fungal Kingdom, J. Heitman, B. J. Howlett, P. W. Crous, E. H. Stukenbrock, T. Y. James, N. A. R. Gow, Eds. (American Society for Microbiology, Washington, DC, 2018), pp. 925–944.

5. M. S. Goettel, J. Eilenberg, T. Glare, in Comprehensive Molecular Insect Science, vol. 6., L. Gilbert, K. Iatrou, S. Gill, Eds. (Elsevier, Boston, 2005), pp. 361–406.

6. D. W. Roberts, R. J. St. Leger, Metarhizium spp., cosmopolitan insect-pathogenic fungi: Mycological aspects. Advances in Applied Microbiology. 54, 1–70 (2004).

7. F. Sánchez-Bayo, K. A. G. Wyckhuys, Worldwide decline of the entomofauna: A review of its drivers. Biological Conservation. 232, 8–27 (2019).

8. D. F. Q. Smith, E. Camacho, R. Thakur, A. J. Barron, Y. Dong, G. Dimopoulos, N. A. Broderick, A. Casadevall, Glyphosate inhibits melanization and increases susceptibility to infection in insects. PLOS Biology. 19, e3001182 (2021).

9. L. Lacey, Manual of techniques in invertebrate pathology (Associated Press, ed. 2, 2012).

10. K. Rydhmer, E. Bick, L. Still, A. Strand, R. Luciano, S. Helmreich, B. Beck, C. Grønne, L. Malmros, K. Poulsen, F. Elb, J. Lemmich, T. Nikolajsen, Automating insect monitoring using unsupervised near-infrared sensors. arXiv. 2108.05435, 16 (2021).

11. M. Brydegaard, S. Jansson, Advances in entomological laser radar. The Journal of Engineering. 2019, 7542–7545 (2019).

12. I. Rigakis, I. Potamitis, N.-A. Tatlas, I. Livadaras, S. Ntalampiras, A Multispectral Backscattered Light Recorder of Insects’ Wingbeats. Electronics. 8, 277 (2019).

13. C. Kirkeby, K. Rydhmer, S. M. Cook, A. Strand, M. T. Torrance, J. L. Swain, J. Prangsma, A. Johnen, M. Jensen, M. Brydegaard, K. Græsbøll, Advances in automatic identification of flying insects using optical sensors and machine learning. Scientific Reports. 11, 1555 (2021).

14. V. Partel, L. Nunes, P. Stansly, Y. Ampatzidis, Automated vision-based system for monitoring Asian citrus psyllid in orchards utilizing artificial intelligence. Computers and Electronics in Agriculture. 162, 328–336 (2019).

15. F. Khamesipour, K. B. Lankarani, B. Honarvar, T. E. Kwenti, A systematic review of human pathogens carried by the housefly (Musca domestica L.). BMC Public Health. 18, 1–15 (2018).

16. C. Elya, T. C. Lok, Q. E. Spencer, H. McCausland, C.C. Martinez, M. Eisen, Robust manipulation of the behavior of Drosophila melanogaster by a fungal pathogen in the laboratory. Elife. 7 (2018) (available at https://doi.org/10.7554/eLife.34414).

17. S. Keller, V. Kalsbeek, J. Eilenberg, Redescription of Entomophthora muscae (Cohn) Fresenius. Sydowia. 51, 197–209 (1999).

18. A. N. Hansen, H. H. De Fine Licht, Logistic growth of the host-specific obligate insect pathogenic fungus Entomophthora muscae in house flies (Musca domestica). Journal of Applied Entomology. 141, 583–586 (2017).

19. L. Malmros, thesis, Lund University, Mathematics (Faculty of Engineering) (2018).

20. T. M. Cooper, R. J. Mockett, B. H. Sohal, R. S. Sohal, W. C. Orr, Effect of caloric restriction on life span of the housefly, Musca domestica. The FASEB Journal. 18, 1591–1593 (2004).

21. S. S. Ragland, R. S. Sohal, Mating behavior, physical activity and aging in the housefly, Musca domestica. Experimental Gerontology. 8, 135–145 (1973).

22. J. Bi, Y.-F. Wang, The effect of the endosymbiont Wolbachia on the behavior of insect hosts. Insect Science. 27, 846–858 (2020).

23. L. J. Cator, P. A. Lynch, A. F. Read, M. B. Thomas, Do malaria parasites manipulate mosquitoes? Trends in Parasitology. 28, 466–470 (2012).

24. A. K. Tallon, M. G. Lorenzo, L. A. Moreira, L. E. M. Villegas, S. R. Hill, R. Ignell, Dengue infection modulates locomotion and host seeking in Aedes aegypti. PLOS Neglected Tropical Diseases. 14, e0008531 (2020).

## Supplementary references

1. K. Rydhmer, E. Bick, L. Still, A. Strand, R. Luciano, S. Helmreich, B. Beck, C. Grønne, L. Malmros, K. Poulsen, F. Elb, J. Lemmich, T. Nikolajsen, Automating insect monitoring using unsupervised near-infrared sensors. 2108.05435, 16 (2021).

2. Chollet, F. & others, Keras (2015). Available at: https://github.com/fchollet/keras.

3. R Core Team, R: A language and environment for statistical computing. R Foundation for Statistical Computing, Vienna, Austria (2014). URL https://www.R-project.org/.

